# Quantitative live-cell imaging of secretion activity reveals dynamic immune responses

**DOI:** 10.1101/2022.02.09.479547

**Authors:** Mai Yamagishi, Kaede Miyata, Takashi Kamatani, Hiroki Kabata, Rie Baba, Yumiko Tanaka, Nobutake Suzuki, Masako Matsusaka, Yasutaka Motomura, Tsuyoshi Kiniwa, Satoshi Koga, Keisuke Goda, Osamu Ohara, Takashi Funatsu, Koichi Fukunaga, Kazuyo Moro, Sotaro Uemura, Yoshitaka Shirasaki

## Abstract

The measurement of cytokine secretions has contributed to the development of immunology; however, new methods that enable highly sensitive and efficient analysis are required for the precise characterisation of dynamic secretion activity when using rare cells or limited human specimens. Here, we report a new technology for quantitative live-cell imaging of secretion activity (qLCI-S), that enables high-throughput and dual-colour detection of prolonged secretion activity at the single-cell level, followed by transcriptome analysis for individual cells based on their phenotype. The power of the qLCI-S was demonstrated by visualising the individual and longitudinal cytokine secretion patterns of group 2 innate lymphoid cells, which comprised <0.01% human peripheral blood mononuclear cells, and identifying their minor subpopulations. This new technology will provide new insights into the spatiotemporal dynamic nature of various secretory functions and the development of fundamental tools for phenotypic drug discovery and regenerative and precision medicine.

## Introduction

Secretion is a fundamental cellular function that is essential for intercellular communication via soluble messengers such as hormones, neurotransmitters, cytokines, growth factors, and extracellular vesicles^1-3^. Cytokine secretion is precisely regulated in a wide range of biological systems such as the immune system, and its dysregulation has been associated with a variety of physiological disorders such as infection, chronic inflammation, allergies, ageing, cancer progression, and autoimmune diseases^4-8^. The detailed characterisation of cytokine secretion is indispensable in our understanding and control of the pathogenesis of these disorders but can be hampered by cellular heterogeneity, which means that even in apparently homogeneous cell populations, each cell is more diverse than we can currently comprehend. Intrinsic fluctuations in cell cycle, history, and activity and stochastic gene expression and regulation cause cellular heterogeneity, and it is eventually reflected in the timing and magnitude of the cytokine secretion^9-11^.

These inevitable cellular heterogeneities mean that bulk evaluations such as enzyme-linked immunosorbent assays and the mass spectrometry of cell populations are not suitable strategies by which to draw definitive conclusions regarding the identity, number, localisation, and individual activities and processes of secretory cells. To address these cellular heterogeneities and to understand complex immune responses analytically and kinetically, it is necessary to develop a new measurement technology that can characterise cytokine secretions in a non-invasive and time-resolved manner with single-cell resolution^12,13^. The high sensitivity and ultimate resolution of the single-cell approach offers considerable advantages when studying the immune response in human clinical specimens in which progress is often limited by the rarity of the target cells^14^. In addition, if this technology can integrate genomic and transcriptomic profiling data for individual cells with the characterisation of their unique secretion activity, the results could be translated into novel and productive scientific research^15-17^.

Currently, enzyme-linked immunospot assays and flow cytometric analysis are well-established techniques applied in both basic research and clinical settings to evaluate cytokine secretion at a single-cell resolution; however, the quantitative nature of the former and the invasive nature of the latter can make their utilisation problematic. To overcome these limitations, several techniques have been developed to enable the non-invasive and quantitative measurement of cytokine secretion at a single-cell resolution using microfabricated nanolitre wells, microfluidic platforms, and water-in-oil droplet technology; these strategies also offer advantages such as high sensitivity, high resolution, high throughput, and the ability to obtain multiparametric measurements^12,18,19^. These excellent techniques, however, have limitations regarding the availability of longitudinal information on secretion activity as they quantify the accumulation of secretion either at a single or a limited number of time points. To correctly evaluate dynamic immune responses, a new technology that characterises the secretion activity of identical cells continuously over a period of days is required.

Fundamental studies have reported techniques that can be used to continuously monitor cytokine secretion at a single-cell resolution^20^. In particular, live-cell imaging of secretion activity (LCI-S), a time-resolved fluorescence immunospot assay performed using total internal reflection fluorescence microscopy (TIRFM), is suitable for a number of reasons. In principle, the detection sensitivity of TIRFM is at the single fluorescent-molecule level, and LCI-S has been demonstrated to have quantitative sensitivity for 2,000 secretory molecules and a temporal resolution time of 1 min^21^. LCI-S can be combined with a variety of imaging techniques that visualise the dynamic features of cells^22-24^. In addition, commercially available antibodies can be used for sandwich immunoassays in LCI-S.

However, there are still technical challenges regarding the use of LCI-S. First, since the signals observed using LCI-S include the kinetics of antibody staining performed in real time, it is necessary to develop an analytical strategy to extract the dynamic secretion activity of living cells^20^. Second, LCI-S needs to be configured for culture conditions that allow for long-term monitoring without impairing cellular activity^25,26^. Third, the LCI-S platform should be stable enough to allow for high-throughput measurements of a statistically comparable numbers of cells. Fourth, the utility of LCI-S can be enhanced if it demonstrates that phenotypically characteristic cells are retrieved from the platform for further analysis.

In this study, we have developed a platform for functional cytology based on quantitative live-cell imaging of secretion activity (qLCI-S platform) by improving the mentioned technical challenges. The performance of qLCI-S has been demonstrated using a novel lymphocyte, group 2 innate lymphoid cells (ILC2), which are known to function through the production of various cytokines^27,28^. ILC2 do not express antigen receptors but are instead activated by the expression of various cytokine receptors. For example, ILC2 maintains moderate proliferative activity in response to interleukin-2 (IL-2). IL-25, produced by tuft cells in epithelial tissues^29^, induces ILC2 to produce type 2 cytokines in the presence of IL-2. IL-33, an alarmin^30^, acts on ILC2 to induce the rapid proliferation and strong production of type 2 cytokines. Because of the continuity of its response and its unparalleled ability for massive cytokine production, ILC2 has a strong influence on the chronicity of allergic diseases such as severe asthma^28,31^, despite its rarity *in vivo*. The rarity of ILC2, especially in human samples, often requires expanded cultures^32^, complicating detailed studies of the secretion response. Therefore, the successful visualisation of cytokine secretion exhibited by freshly isolated human ILC2 as well as the identification and characterisation of rare and heterogeneous subpopulations associated with comprehensive gene expression analysis, clearly demonstrate the power and utility of the qLCI-S platform.

## Results

### Establishment of an LCI-S quantitative analysis

LCI-S is performed on a glass/resin hybrid culture dish with a microfabricated nanolitre well array (Fig. 1a and 1b) and employs a time-resolved fluorescence immunoassay on the bottom surface of each well, which can be visualised using TIRFM. Previously, LCI-S used a laser source for TIRFM^33^, which impaired quantification by illuminating undesirable interference patterns due to coherence. To significantly improve the optical stability of the platform, we devised a fabrication process for an advanced microstructure of the nanolitre well array, using a layer of amorphous fluorocarbon (AF) polymer for refractive index coordination, which was sandwiched with a glass substrate and microfabricated polydimethylsiloxane (PDMS). This structure allowed the use of TIRF illumination with incoherent light-emitting diodes (LED-TIRF)^34,35^ in the LCI-S platform, ensuring the detection of sufficient fluorescence signal from the target at small incident light angles, while reducing the background signal from leakage light (Fig. 1c, d). We evaluated the effect of an optically stable LED-TIRF illuminator on the quality of the LCI-S measurement signal. The same wells were imaged with both LED-TIRF illuminator and Laser-TIRF illuminator and compared to assess the uniformity of the fluorescence intensity (Fig. 1e). Comparisons of brightness variation of the illumination patterns within the wells, the well-to-well variation in mean fluorescence intensity, and the temporal fluctuations of the mean fluorescence intensity in individual wells during time-lapse imaging showed significantly lower CVs with the LED-TIRF illuminator than the Laser-TIRF illuminator (Fig. 1f). Thus, the LCI-S platform successfully improved the measurement stability sufficient for quantitative signal analysis by developing an optically advanced nanolitre well array structure that allowed the installation of LED-TIRF illuminator, which is more stable than Laser-TIRF illuminator.

**Figure 1.**
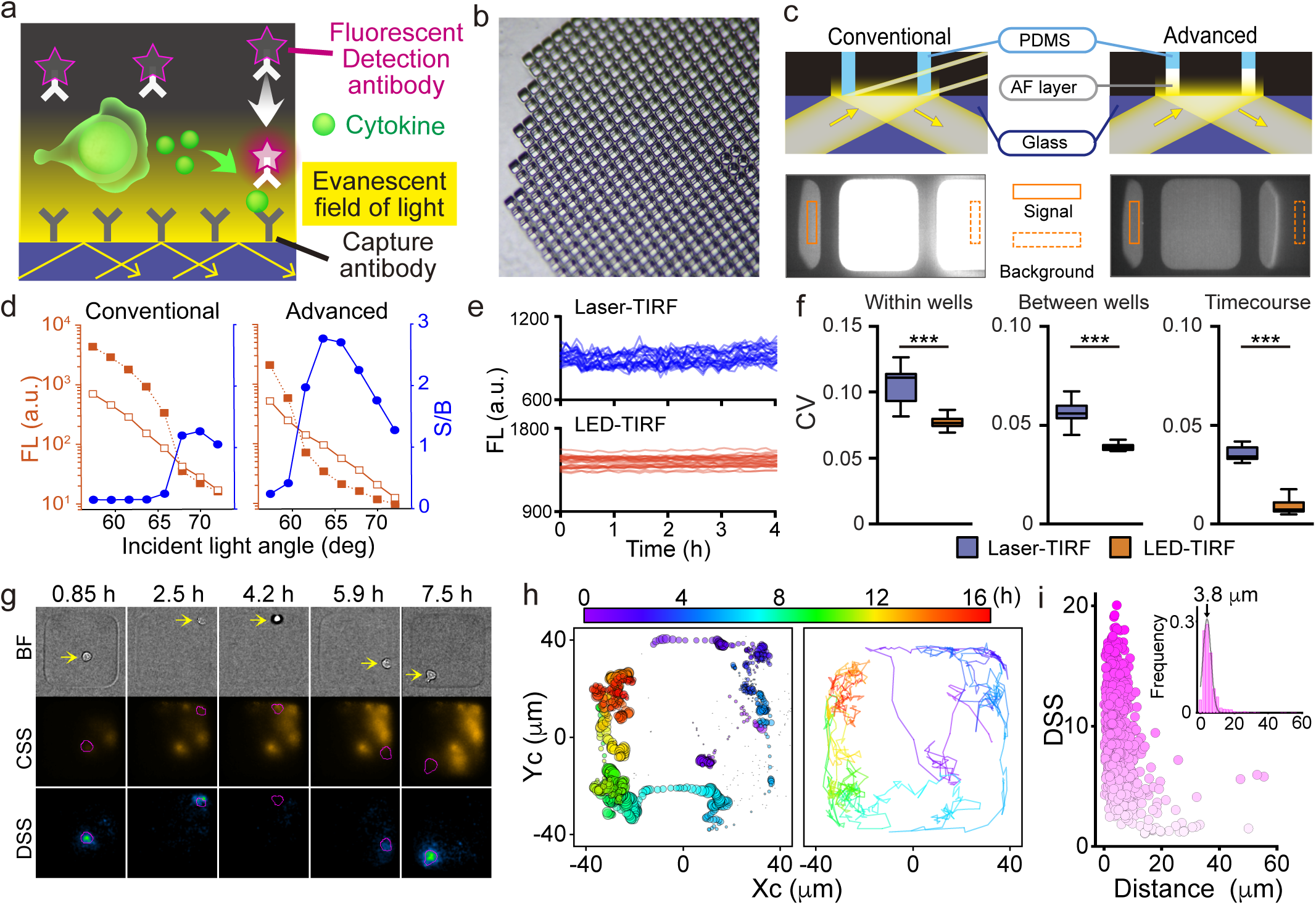
Establishment of the qLCI-S platform. **(a)** Scheme for the time-resolved fluorescence immunospot assay on total internal reflection fluorescence microscopy (TIRFM) for live-cell imaging of secretion activity (LCI-S). **(b)** Overview of the microfabricated nanolitre well array structure. Illustration of the qLCI-S platforms optical stability. The advanced microfabricated well structure consisted of a bottom glass substrate and a nanolitre well array composed of polydimethylpolysiloxane (PDMS) with a refractive index adjusting layer of amorphous fluorocarbon polymer (AF layer) sandwiched in between. High background signals are generated by stray light from the interface between the glass and PDMS in the conventional structure. The AF layer inserted into the interface between the glass and PDMS prevents the generation of stray light. **(d)** Effects of the AF layer on improving the ratio of signal-to-background noise. Changes in the signal intensity (red open squares, mean intensity of the area with no stray light overlapping, outlined by solid lines and indicated as “Signal” in (c)) and background noise (red filled squares; mean intensity of the area illuminated only with stray light, outlined by dash lines and indicated as “Background” in (c)) along with the angles of the incident excitation light without (left) or with (right) the AF layer were plotted. The signal-to-noise ratio (blue filled circles) was calculated from these values. **(e)** Comparison of the temporal stability of fluorescent signals with the laser-TIRF illuminator and the LED-TIRF illuminator. The traces of mean fluorescent intensity of individual wells illuminated with the laser-TIRF (upper panel, blue lines) and LED-TIRF (lower panel, red lines) illuminator were shown. **(f)** Comparison of the spatiotemporal variation of the TIRF signals. Left panel shows the coefficient of variation (CV) of intensities within the wells at each time points, which reflected the intensity variation of interference pattern of the laser or LED light. Middle panel shows the CV between wells at each time points. Right panel shows the CV of temporal fluctuation of each well. Differences were assessed using the Mann–Whitney *U* test (*** p < 0.001). **(g)** Bright-field images of a mouse innate lymphoid cell (mILC2; top, BF), cumulative secretion signal (CSS) images of IL-5 (middle, CSS), and temporally deconvoluted secretion signal images [bottom, deconvoluted secretion signal (DSS)]. Images at representative points in time from the data acquired and analysed every minute are shown. **(h)** Left: The 2D Gaussian distributions fitted to the DSS signal at each time point are indicated by the position (centre), size (height), and colour (time) of the circles. Right: Movement of the geometric centre of ILC2 at each time point is shown as a trajectory drawn in time-coded colours. **(i)** Relationship between the intensity of the DSS and the distance of its centre from the cell; the higher the intensity, the closer the central position of the DSS to the cell location. The inset histogram shows the distribution of the distances with a peak at 3.8 μm.

Next, we verified that the novel analytical qLCI-S strategy, which temporally deconvoluted antibody staining kinetics from a series of secretion signal images, would allow us to extract dynamic secretion activity in a spatiotemporally-resolved manner. For the demonstration, we measured the IL-5 secretion of ILC2 from mouse adipose tissue (mILC2) stimulated with IL-33, and calculated the dynamic IL-5 secretion activity. The deconvoluted secretion signal (DSS) was calculated from the cumulative secretion signal (CSS) image of IL-5 using the method described in the Materials and Methods, which utilised reference data for the staining kinetics of the IL-5 antibody against the recombinant IL-5 protein under the same measurement conditions (Fig. 1g, Supplementary Video 1). From a CSS image that showed the signal overlapping throughout the nanolitre well over time, we generated a DSS image showing how the local signal distribution moved over time. To clarify the spatial relationship between the localised signal and cell, we fitted the signal distribution in the DSS image at each time point using a Gaussian distribution function (Fig. 1h) and measured the distance between the centre of the signal distribution and the geometric centre of the cell position. We found that both centres were co-localised at a close range (mode distance 3.8 μm) (Fig. 1i). This clearly demonstrated that qLCI-S could be used to visualise dynamic cellular secretion activity with a high spatiotemporal resolution.

Based on these results, we used the observation dish (TIRF-dish), which featured an advanced nanolitre well array with glass/AF/PDMS layers compatible for stable LED-TIRF illumination, for subsequent experiments.

### Configuration of qLCI-S for long-term monitoring

To obtain qLCI-S measurements over a period of several days without impairing cellular activity, we assessed the impacts of the single-cell culture environment and phototoxicity by fluorescence imaging. For sensitive and quantitative detection of the cytokine secretion, isolating the target cell in a closed nanolitre space (enclosed cultivation) is theoretically effective. However, the resources required by the cells are limited in the small volume of the nanolitre space. The effects of rapid environmental changes in the limited space associated with cell metabolism must also be considered^36^. Previously, a volume-dependent decrease in antibody production was reported in enclosed cells^37^. In contrast, isolating the target cell in an open-ended nanolitre space (semi-enclosed cultivation) maintains a constant environment, similar to traditional *in vitro* culture procedures. In addition, a study on the simulation in open-ended nanolitre space in an immunospot assay showed a correlative signal outcome with the secretion activity of the cells^38^. Therefore, we evaluated whether enclosed or semi-enclosed cultivation was suitable for qLCI-S by comparing the detection sensitivity and persistence of secretion activity of mILC2s (Fig. 2a). We did not observe any significant differences in the CSS of IL-5 and the rate of secretion-positive cells between the enclosed and semi-enclosed cultivation after 3 h. Conversely, after 24 h, the CSS was significantly lower under enclosed cultivation than under semi-enclosed cultivation (Fig. 2b, c; p < 0.001). The inhibition of secretion activity under enclosed cultivation was also remarkably reflected in the rate of secretion-positive cells, as they were reduced to approximately half (Fig. 2d). In addition, morphological comparison of the secreting ILC2s showed that the semi-enclosed cultures were hypertrophied whereas in comparison the enclosed cultures were transformed but smaller.

**Figure 2.**
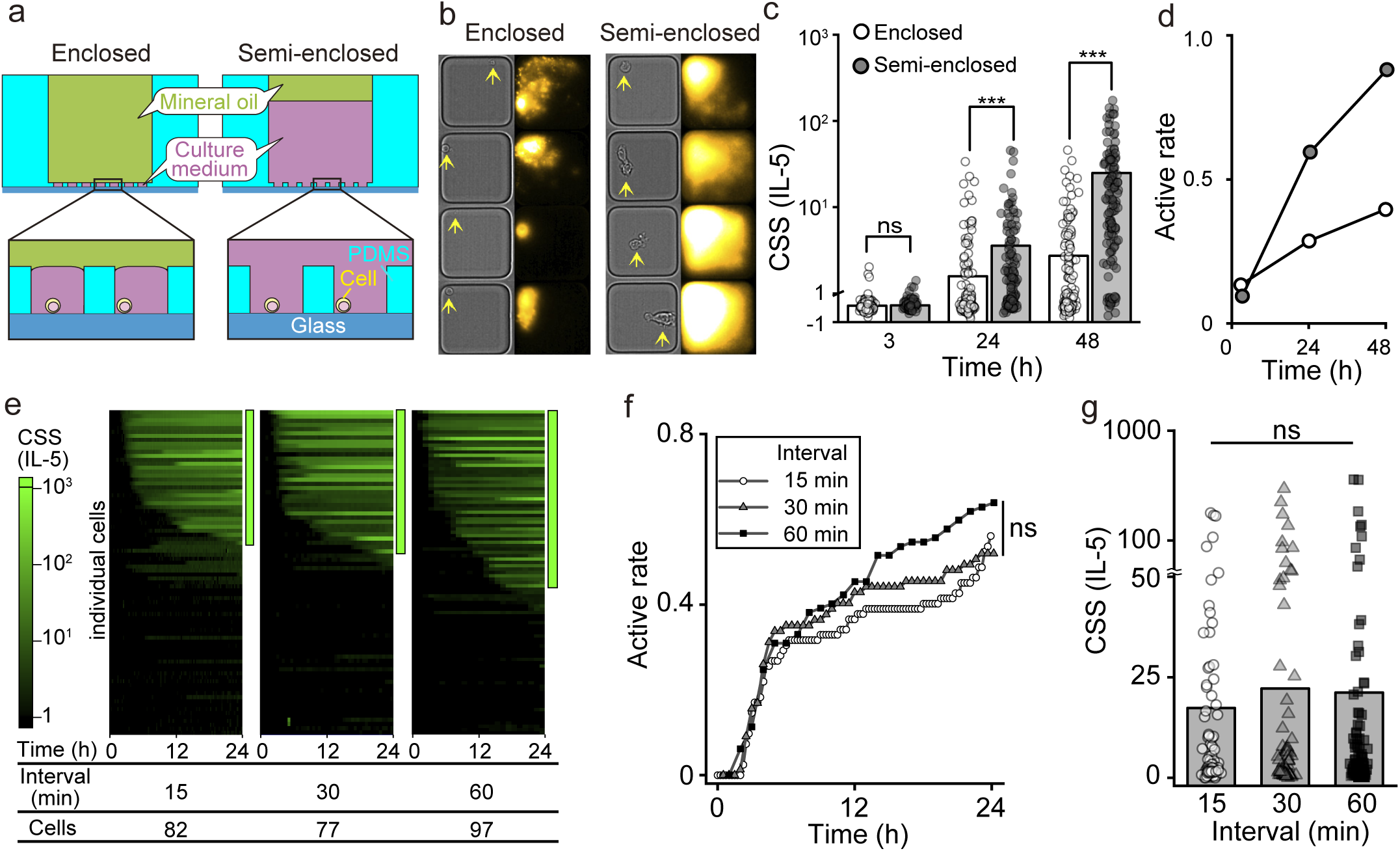
Evaluation of the qLCI-S platform. **(a)** Schema of the experimental setup for the single-cell cultivation with the nanolitre well array. For “enclosed cultivation”, cells (yellow) were sealed in nanolitre wells with mineral oil (green) along with a limited volume of culture medium (purple); for semi-enclosed cultivation, cells were deposited in open-ended nanolitre wells surrounded by a large pool of culture medium. **(b-d)** Comparison of IL-5 secretion activity between the enclosed and semi-enclosed cultivation. (b) The representative images of the secretion activity under the two types of cultivation 48 h after isolation. Yellow arrows indicate the position of the cells. Comparison of the cumulative secretion signal (CSS) from individual cells (c) and the activation rate of mILC2s (d) in each cultivation state. *** p < 0.001 or ns p > 0.05; non-parametric Mann–Whitney test between the two cultivation states. **(e)** Chronological heatmap of the CSS of individual cells tracked by qLCI-S for the indicated number of cells with the indicated time-lapse intervals. The colour reflects the CSS intensity. Each row reflects the dynamic secretion activity of an individual cell. The colour scale bar shows the intensity of IL-5 secretion signals. Secretory positive cells are indicated by green lines on the right side of the chronological heatmap. **(f)** Rate of activation of mILC2s at different time-lapse intervals. Differences were tested using the log-rank test. **(g)** Maximum CSS of each cell with different time-lapse intervals. Differences were assessed using the Mann–Whitney *U* test.

Next, we evaluated the effects of phototoxicity during the qLCI-S observations. Assuming that the frequency of shooting is proportional to the phototoxicity, the IL-5 secretion activity of mILC2s was measured at intervals of 15, 30, and 60 min and the activities under each condition were statistically compared. The results showed no significant differences in the rate of secretion-positive cells (Fig. 2e, f) or the CSS (Fig. 2g).

Based on these results, we concluded that qLCI-S can be effectively used to follow the actual secretion activity of living cells for several days with semi-enclosed cultivation and a one-hour time-lapse.

### Statistical comparison of secretion activity using the qLCI-S platform

The qLCI-S platform was evaluated for its ability to provide statistically comparable high-throughput measurements using a TIRF-dish consisting of four chambers with 996 wells at the bottom of each chamber (Fig. 3a). In this demonstration, the production of cytokines IL-5 and IL-13, which are typical of mILC2, were measured simultaneously using a mixture of immobilised capture antibodies and differentially coloured fluorescent detection antibodies (Fig. 3b). The equivalence between the four chambers was confirmed when the same experiment was performed in parallel in all four chambers of the TIRF-dish, yielding four observations of equivalent cytokine secretion activity for the mILC2s (Supplementary Fig. 1). Next, we compared the stimulus-dependent cytokine production responses of mILC2 using four types of stimuli known to elicit different responses: IL-2, IL-2/IL-25, IL-33, and IL-2/IL-33 (Fig. 3c). Most of the mILC2s activated with IL-2/L-33 stimulation produced either IL-5 or IL-13, 12 h after stimulation (rate: 0.67). The rate for double-positive cells was 0.57, and the rates for IL-5 and IL-13 single-positive cells were 0.06 and 0.04, respectively (Fig. 3d). The intensities of the secreted IL-5 and IL-13 were correlated on a logarithmic scale within cells, and significantly differed by more than two orders of magnitude between cells (Fig. 3e). Although active mILC2s (rate: approximately 0.7) started secretion within 5 h of stimulation, the starting points varied widely from 1–12 h after stimulation for IL-2/IL-33 (Fig. 3f). The start of secretion for the two cytokines was coincidental but not always synchronous (Fig. 3g).

**Figure 3.**
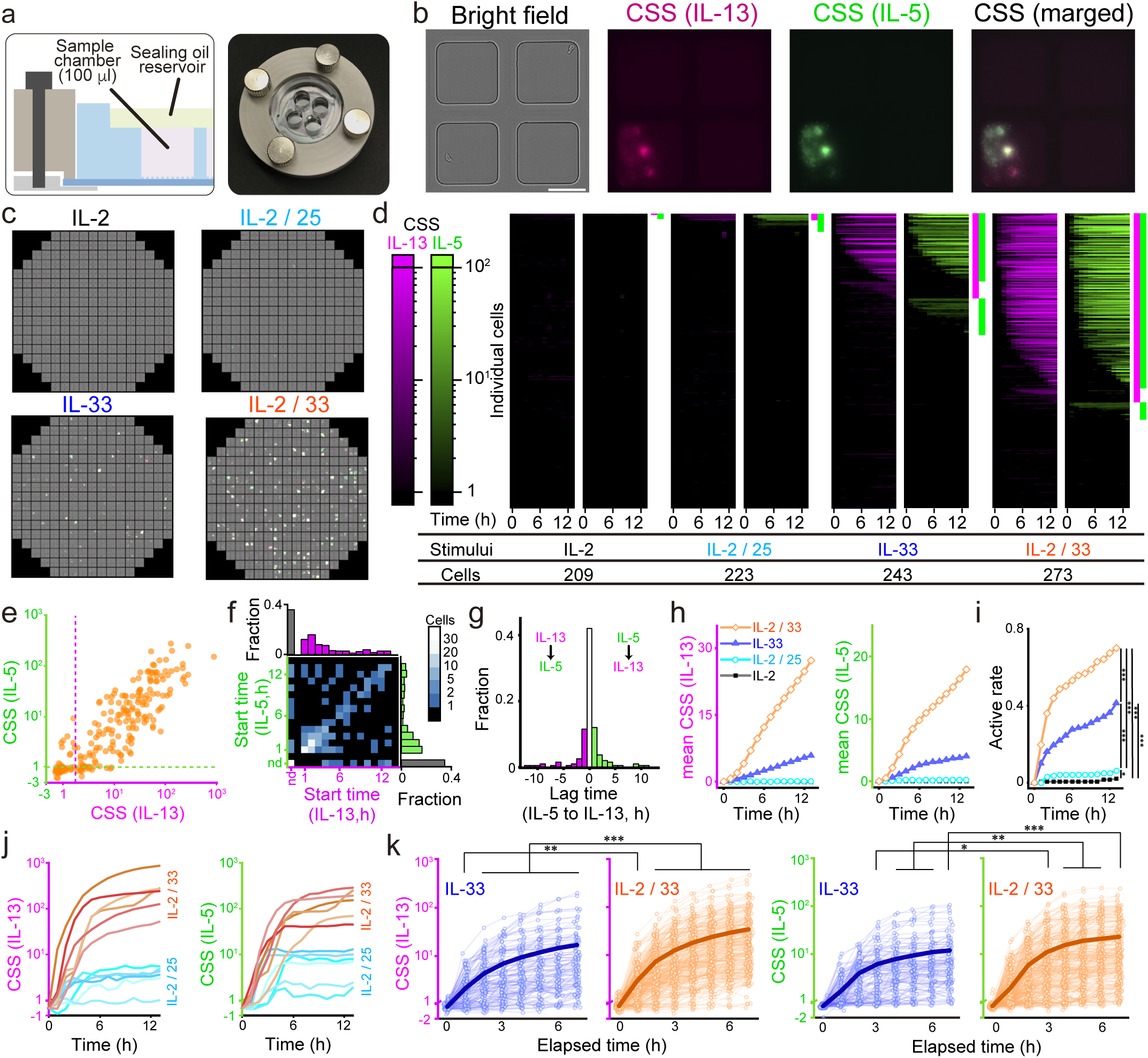
Statistical comparison of the secretion activity using the qLCI-S platform. **(a)** Overview of the quadripartite TIRF-dish with four chambers with open-ended nanolitre well arrays and a space for oil-sealing to cover them. The dish is held with a home-made jig for installation on the microscope. **(b, c)** Secretion activity of the cells in the nanolitre wells was tracked using LCI-S. (b) Representative example of LCI-S data obtained from a single field of view. Bright field (left), IL-13 signal (magenta), IL-5 signal (green), and IL-5/IL-13 superimposed images (merged) are shown. (c) Overall view of responses from individual mILC2s stimulated with different stimuli. Each of the 249 fields of view was arranged according to its position on the TIRF-dish. **(d)** The chronological heatmap of CSS from the individual mILC2s under different stimuli. Each row reflects the dynamic activity of individual mILC2s for the total number of single cells indicated for each stimulus, whereas the column indicates the time series. The colour scale bars show the intensity of CSS (IL-13: magenta, IL-5: green). Time-lapse data were obtained every 1 h. **(e)** Comparison of the CSS intensity of the two cytokines in each mILC2. Each axis corresponds to a cytokine species while each point corresponds to a cell. The dashed line on each axis indicates the detection limit. **(f, g)** Comparison at the start of secretion. (f) Correlation of the time from stimulation to the start of secretion between cytokines; cells without detectable secretion after 12 h of observation were assigned to the origin of each axis in black colour as not detectable (n.d.). The number of cells in each bin are displayed using a heat map according to the colour bar shown on the right. (g) Differences at the start of secretion between the two cytokines in each cell; 40 % of the cells started to secrete at the same time, but some were preceded by IL-13 (magenta) and some by IL-5 (green). **(h)** Comparison of the mean CSS of ILC2s under different stimuli. The mean CSS was calculated from the total sum of the CSS divided by the total number of analysed single ILC2s, as comparable to the bulk secretion assay. **(i)** Comparison of the rate of activation of the ILC2s under different stimuli. ILC2s with a CSS of IL-5 or IL-13 over each threshold based on the CSS of empty wells are shown. Significant differences were assessed for all group combinations using the Benjamini– Hochberg corrected log-rank multiple test (*** p < 0.001, * p < 0.05 with q < 0.01). **(j)** Comparison of secretion activity between IL-2/25 (blue) and IL-2/33 (orange) stimuli: the traces of top six in CSS with each stimulus are shown for IL-13 and IL-5 secretion. **(k)** Comparison of the secretion activity of individual cells stimulated with IL-33 in the absence (blue) or presence (orange) of IL-2. The evolution of the signal from the start of secretion is aligned. The thin lines show the trace of each cell, and the thick lines show their mean. Differences were assessed using the Mann–Whitney *U* test (*** p < 0.001, ** p < 0.01, * p < 0.05).

We then performed a comparative analysis among the different stimuli. First, we compared the ensemble averages of the secretion signals, which mimic the conventional bulk measurements of the cell population, to determine whether the results obtained are in line with previous findings (Fig. 3h). mILC2 responded most strongly to IL-2/IL-33 stimulation, followed by weaker responses to IL-33, IL-2/IL-25, and IL-2, which agreed with previous studies^39^. We further compared the response to each stimulus in terms of the secretion activity at the single-cell level. Particularly prominent was the difference in the rate of secretion-positive mILC2s in response to each stimulus (Fig. 3i, p < 0.001), which proved comparable to the ensemble mean of the secretion signals described above. Next, we focused on the differences in the trace patterns of the secretion activity of individual cells for each stimulus. Comparison of representative traces of highly secreting cells stimulated with IL-2/IL-25 or IL-2/IL-33 revealed a characteristic difference: stimulation with the former resulted in a small amount of transient secretion whereas stimulation with the latter resulted in a large amount of continuous secretion (Fig. 3j). Although IL-2 alone could not induce cytokine production (Fig. 3c, d, h, i), co-stimulation with IL-33 enhanced the intensity of the secretion activity of individual mILC2s (Fig. 3k, p < 0.001) as well as the positive rate of secretion. These results demonstrated that the qLCI-S platform provides a high-throughput measurement that allows for statistical comparison of the detailed differences in secretion activity in a small number of cells of the same batch.

### Influential minor ILC2 regulates the entire innate type 2 secretion response

After evaluating the mILC2 grown in the *in vitro* expansion culture, we performed the qLCI-S analysis of ILC2 from human peripheral blood (hILC2). As the fraction of hILC2s is typically small and insufficient, conventional methods require prior expansion cultivation of hILC2s to assess cytokine secretion^40^. We investigated whether the qLCI-S platform could track the secretion activity of the freshly purified hILC2s (Supplementary Fig. 2) without expansion cultivation, which is close to their *in vivo* state with the potential to provide insights into the diversity of the immune response depending on the donors.

First, we focused on hILC2s that exhibited high amounts of IL-5 and IL-13 secretion upon stimulation with IL-2 and IL-33 and observed how these cells developed their secretory functions over 5 days (Fig. 4a, Supplementary Video 2). As the secretion activity progressed, hILC2s transformed from a small spherical shape to an enlarged shape with a tail and fin-like structures. In most cases, IL-5 and IL-13 secretion activities were synchronised, but their intensity fluctuated at around 20 h (Fig. 4b). The continuously secreting ILC2s underwent cyclic proliferation and repeatedly attached and dissociated into a single aggregate. In wells where large amounts of cytokines were produced along with repeated proliferation, the increase of CSS was affected by the capacity of the capture antibody (Fig 4b). Therefore, the analysis of the secretion activity of hILC2 according to the intensity of CSS was performed up to day 3 after stimulation, when the activated hILC2 did not show more than three divisions.

**Figure 4.**
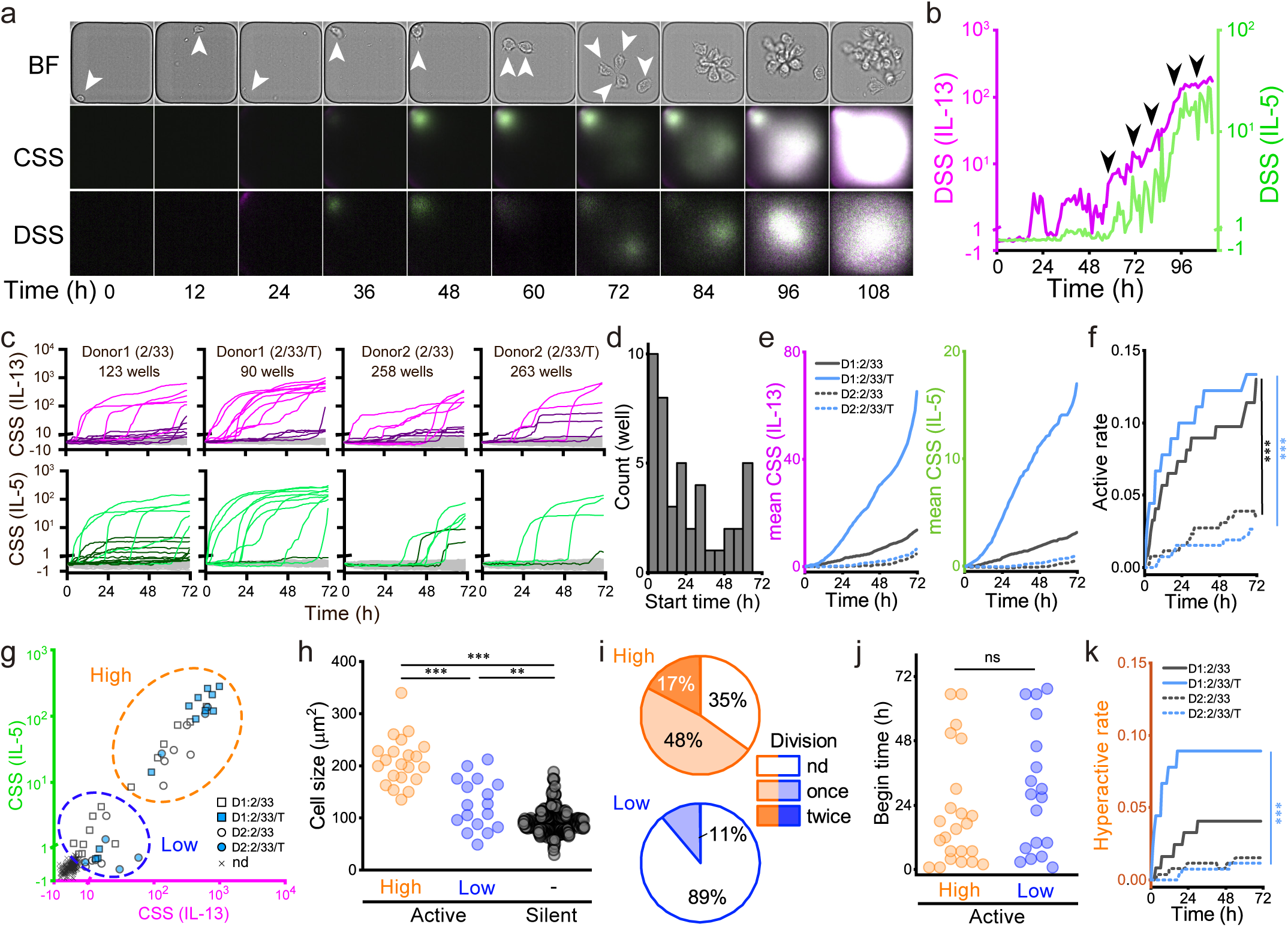
Tracking the type 2 cytokine secretion activity of hILC2s. **(a, b)** Tracking of hILC2 secretion activity using the qLCI-S platform. Bright-field images of an hILC2 (top, BF); CSS and DSS images in the superimposed colour of IL-5 (green) and IL-13 (magenta). Data are shown for every 12 h. Arrowheads indicate the position of hILC2 until the second division. **(c)** Chronological trace of the secretion activity of hILC2s for 3 d. CSS of each well summarised for each donor (D1: donor 1 or D2: donor 2) and each stimulus (2/33: IL-2 and IL-33 stimulation, 2/33/T: IL-2, IL-33, and TSLP stimulation). Each line reflects individual wells containing a single cell or several cells, as estimated by the AI-supported cell counter. The total number of cell-containing wells is shown for each sample index. Bright, dark, and light-grey lines indicate hyperactive, slightly active, and silent ILC2s, respectively. **(d)** Distribution at the start of secretion in individual hILC2s. The start of secretion for all cells detected in (c) was made into a histogram, regardless of the stimulus type or donors. **(e, f)** Statistical comparison of the secretion activity between the two donors with or without TSLP. Comparison between donors and stimulations (solid lines: donor 1, dashed lines: donor 2, black: IL-2/IL-33 stimulation, blue: IL-2/IL-33/TSLP stimulation). (e) Comparison of the mean CSS (IL-5: green, left; IL-13: magenta, right). (f) Comparison of the rate of activation of ILC2s. Differences were tested using a Benjamini–Hochberg corrected log-rank multiple test (*** p < 0.001 with q < 0.01). **(g)** Classification of activated ILC2 based on their levels of secretion signal. Comparison of the maximum CSS over time for IL-5 and IL-13 in each trace shown in (b), squares and circles indicate traces from active ILC2 of D1 and D2, respectively. Blue and white colours indicate the absence and presence of TSLP, respectively. A cross indicates traces below the threshold. The detectable trace data were divided into two groups using k-means clustering: hyperactive (high) and slightly active (low). **(h)** Comparison of the cell sizes among the 3 groups with different secretion activities. The average area of the cells obtained from the bright-field image observations of the last six time points in a 72-h period was used as a size parameter. Cells contained within each well were classified based on the CSS levels detected from each well, as shown in (c); when wells contained more than a single cell, their average area was adopted as a representative value. Differences were assessed using the Mann– Whitney *U* test (*** p < 0.001, ** p < 0.01). **(i)** Comparison of the division activity between the two groups with different secretion activities. The number of divisions in the activated ILC2s was counted for 72 h using an AI-supported cell counter followed by visual verification. **(j)** Start of secretion comparisons for the two groups in active hILC2 with different secretion activities. There is no difference in the start of secretion between the hyperactive and slightly active hILC2s. Differences were assessed using the Mann– Whitney *U* test. **(k)** Comparison of the rate of activation of ILC2 resulting in hyperactivity. Differences were tested using a Benjamini–Hochberg corrected log-rank multiple test (*** p < 0.001).

Next, to investigate how individual differences in type 2 immune responses can be described using the single-cell characteristics of hILC2s, we statistically compared the secretion activity of hILC2 from different donors stimulated by IL-2/IL-33 with or without thymic stromal lymphopoietin (TSLP), which is an effector of ILC2 survival ^41^ (Fig. 4c). Throughout the 3 days of continuous observation, only a fraction of the hILC2 showed IL-5 and IL-13 secretion activity (Fig. 4c). The time course for secretion activity from each activated hILC2 was individualised; hence, the start of secretion was widely distributed, ranging anywhere from hours to days after stimulation (Fig. 4d). A statistical comparison of the secretion activity of the hILC2s between the donors showed contrasting effects for the TSLP; donor 1 showed a TSLP-dependent increase in the mean secretion amount, which was not observed in donor 2 (Fig. 4e). Conversely, secretion-positive cells, which showed a significant difference between donors (Fig. 4f, p < 0.001), were unaffected by TSLP stimulation in either donor (Fig. 4f). This indicates that only hILC2 from donor 1 was affected by TSLP when there was an increase in the intensity of the secretion activity by individual cells.

Next, we carefully compared the secretion activity tracks for the individual hILC2s and identified a pattern in the intensity differences and persistence of the secretion activity, similar to that observed in the mILC2 stimulated with IL-25 and IL-33. A classification depending on the CSS of individual hILC2s at 72 h after stimulation revealed the existence of three groups: hyperactive (high), slightly active (low), and silent (Fig. 4g). In particular, the hyperactive hILC2s had other striking features such as morphological enlargement (Fig. 4h, p < 0.001) and high proliferative activity (Fig. 4i). These differences in the secretory function may not be attributed to differences in response duration, as the distribution when secretion first started was not different between the slightly active and hyperactive hILC2s (Fig. 4j). Focusing on the persistence of the secretion activity, the hyperactive hILC2s tended to prolong the secretion activity; whereas, the slightly active hILC2s tended to cease cytokine production in 10–30 h (Fig. 4c). We statistically compared the secretion activity of the hyperactive hILC2s between donors. The results showed that the TSLP-stimulated donor 1-derived hILC2s had a significantly higher rate of hyperactive cells than those under other conditions, and the trend in the rates of the hyperactive cells was similar to that of the mean secretion amount (Fig. 4k, p < 0.001). These results suggested that TSLP enhanced the hyperactivation of hILC2s, resulting in an increase in the total secretion amount as well as the number of hILC2s at the population level, although the strength of the effect on hILC2 varied among donors.

In summary, qLCI-S revealed that the type 2 cytokine secretion for IL-33/TSLP stimulation that was expected in freshly isolated hILC2 was mediated only by a small but hyperactive subpopulation that had not previously been identified, and this could also be referred to as an influential minority.

### Comparison of gene expression between hyperactive and silent ILC2s

The hILC2 population was classified by qLCI-S into hyperactive, slightly active, and silent subpopulations according to the secretory function reflected in the phenotypic differences in secretion and proliferation activities and morphological changes. The comprehensive gene expression information underlying the differences in the secretory function promised to deepen our understanding of the regulatory mechanisms of hILC2 activation. We, therefore, tested the feasibility of cell selection and recovery based on the observed secretion activity and its application to single-cell transcriptome analysis. After 5-days of qLCI-S for the ILC2s stimulated with IL-2/IL-33/TSLPs, hyperactive and silent phenotypes were harvested as individual clones and single cells, respectively (Fig. 5a), and applied to RNA-Seq analysis. Comparison of the mean transcripts per million (TPM) values for the hyperactive and silent phenotypes revealed the specific gene expression for each phenotype, e.g. the gene expression of IL5 and IL13 was significantly higher in the hyperactive group (Fig. 5b and see the explanation of “DEG” in Materials and Methods). Conversely, we confirmed that the expression of a series of genes corresponding to the isolation markers of hILC2 used in this study was common in both groups (Supplementary Fig. 3a), ensuring that both could be regarded as hILC2s with completely different phenotypes.

**Figure 5.**
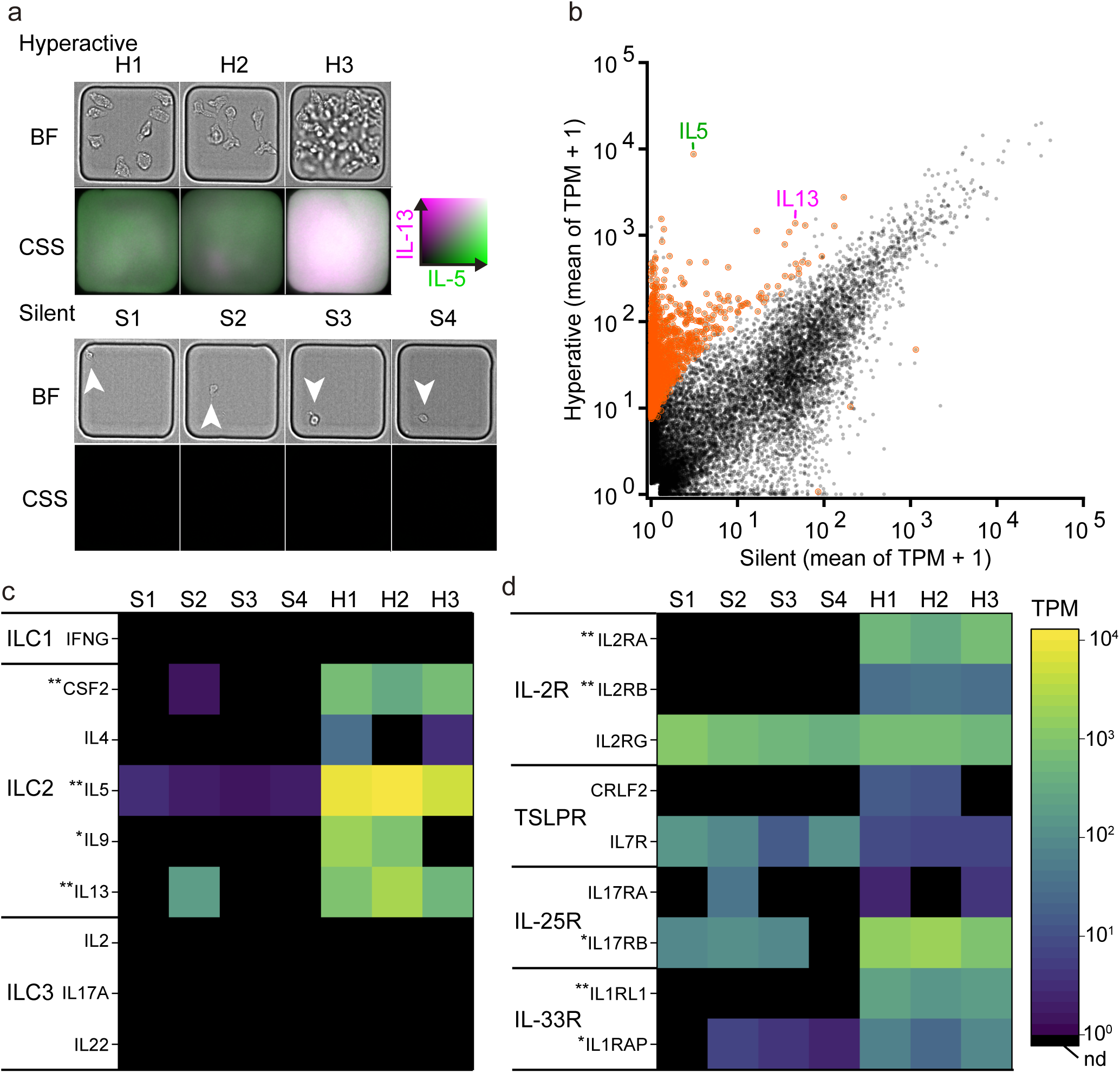
Functional transcriptome analysis of hILC2. **(a)** Images showing cell morphology (upper) and cumulative secretion activities (lower, IL-13: magenta, IL-5: green) of three hyperactive ILC2 clones (H1–H3) and four silent ILC2s with undetectable secretions (S1–S4) collected for RNA-Seq. **(b)** Scatter plot of mean transcripts per million (TPM) of the hyperactive (high) and silent ILC2s. DEGs with a p-value < 0.01 and false discovery rate (FDR) < 0.01 are coloured in red. **(c)** Heatmap showing the expression levels of the characteristic cytokine genes of ILCs. **(d)** Heatmap showing the gene expression of the receptors required to activate the stimulation of ILC2s. Differences were tested using a ROTS (*** p < 0.001, ** p < 0.01 with FDR < 0.05).

To explore the origin of the different phenotypes reflected in the transcriptome, we investigated the genes differentially expressed between the hyperactive and silent ILC2s. For example, CSF2^42^ and IL9^42,43^ were predominantly expressed in hyperactive ILC2s as well as IL5 and IL13 (Fig. 5c, p < 0.05 for IL9 and p < 0.01 for others). IL4 was slightly expressed in hyperactive ILC2s (Fig. 5c), whereas ILC1 and ILC3 cytokine characteristics were not expressed in either the hyperactive or silent ILC2s. Furthermore, among the other gene characteristics of hILC2, the transcription factor RORA^43^ and the immune checkpoint protein ICOS^43^ tended to be highly expressed in hyperactive ILC2s (Supplementary Fig. 3b). These results indicated that silent ILC2s were not activated, even at the transcriptional level. Based on this indication, the silent ILC2s were in a cellular state that was insensitive to stimuli. Therefore, we focused on gene expression of the receptors for hILC2-activating stimuli and compared the results between hyperactive and silent ILC2s and found that the silent ILC2 barely expressed a subset of heteroreceptor genes for IL-2 and IL-33 activating stimuli (Fig. 5d, p < 0.01). These results indicated that the silent hILC2s found in qLCI-S were functionally branched hILC2s that are insensitive to stimuli and differed from the hyperactive hILC2s at the gene expression level.

The findings from this functional transcriptome analysis demonstrated the scalability of qLCI-S in combination with various subsequent approaches for cellular characterisation.

## Discussion

In this investigation, we have established the qLCI-S platform, which integrates high-throughput, long-term, and dual-colour quantitative tracking of secretion activity with transcriptomic analysis of the identified phenotypic cells.

The first achievement in this study was assuring that the time-resolved signal of the fluorescence immunospot assay on the total reflection fluorescence microscope, reflected the secretion activity of the cells. We demonstrated the imaging of IL-5 secretion activity for mouse ILC2 as a typical example of secretion through the classical secretory pathway. The improved optical stability reduced the noise in the LCI-S signal and consequently, explicitly showed that the LCI-S quantitative analysis with the high spatiotemporal resolution of qLCI-S enabled to identify the dynamic cytokine secretion of mILC2, which, previously, had only been speculated.

The second achievement in this study was determining that quantitative tracking of the secretion activity over 3 days could be achieved with a throughput that was sufficiently high as to allow for statistical analysis. These measurement specifications for cytokine secretion were not previously reported with single-cell techniques^20^. In addition, the ability of the qLCI-S platform to evaluate the secretion activity of two cytokines simultaneously and in parallel under four conditions maximised its advantage when working with human specimens, which have limited availability and the possibility of inter-assay variation under sample conditions. In fact, by taking advantage of this, we uncovered the stochastic nature of the start of cytokine secretion exhibited by mouse and human ILC2, which is broadly distributed in the temporal direction over 1 to 12 h (most frequently 2 h) for mouse and 1 to over 72 h (most frequently 3 to 6 h). The stochastic nature of immune activation, in which the rate of activation in the population varies with stimulus concentration, has been reported in previous studies^9,44^. However, in this study, ILC2s were provided with a sufficient concentration of stimulus, so that they were not depleted during long-term cultivation. Therefore, the stochastic nature of the activation exhibited by ILC2 was assumed to be controlled by an endogenous molecular mechanism that dispersed the timing of activation.

In this study, we also discovered the existence of a minor subpopulation of hyperactive hILC2s, which are highly influential due to their dominant secretory and proliferative activities. There were individual differences in the rate of the hyperactive hILC2s and the impact of the rate by TSLP. These individual differences in the cytokine secretion of hILC2s may be associated with pathological conditions such as asthma severity and allergic constitution. Thus, these differences must be further investigated in a large number of individuals.

The third achievement of this study was the implementation of functional transcriptome analysis based on the phenotyping of secretory functions. The stimuli receptors, such as IL-2, IL-33, and TSLP, were highly expressed in hyperactive hILC2s and negatively regulated in silent hILC2. Considering that freshly purified ILC2s from human peripheral blood have low expression of the IL-33 receptor^45^, functional fate determination of ILC2 activation and maintenance of hyperactivity presumably involves intracellular molecular mechanisms that control the promotion and maintenance of stimuli receptor expression. Comparing the dynamics of stimuli receptor expression and secretion activities would shed light on the detailed molecular mechanisms of ILC2 activation.

The qLCI-S also has some technical challenges that need to be addressed. The first is the limitations in the trade-off between throughput and temporal resolution. The throughput of qLCI-S depends on the size of the field of view of the objective lens for total reflection fluorescence microscopy and on the speed of scanning of the microscope system, which was approximately 4,000 shots/h in this study. Innovations such as the widening of the field of view, a faster microscope system control, and simultaneous multi-colour imaging capabilities will improve the throughput and temporal resolution of qLCI-S. Second is the evacuation of cells from open-ended nanolitre wells. In this study, the high migration ability of mouse ILC2s made it challenging to evaluate long-term tracking over several days. Innovations in the structure of nanolitre wells are expected to interfere with the escape of cells from the wells. The third challenge is the multiplexing of measurement targets. Improvements are expected to be made by increasing the variety of colours for different detection antibodies and the variety of capture antibodies via micropatterning of the solid phase surface.

Despite the limitations described above, qLCI-S has practical advantages such as the simplicity of the measurement process, the availability of commercially available antibodies, and the ease of integration with existing live-cell imaging techniques. QLCI-S may have applications, like visualising the secretion activities of various intercellular messengers such as cytokines, neurotransmitters, and extracellular vesicles and of dynamic cell–cell interactions. Further innovations in qLCI-S and the development of applied research are expected to provide new perspectives in areas such as the elucidation of molecular mechanisms in diverse diseases, phenotypic drug discovery, provision of resources for regenerative medicine, and criteria for precision medicine.

## Methods

### Reagents

Antibody pairs used for the fluorescence immunospot assays of mouse IL-5 (monoclonal capture antibody MAB405 and monoclonal detection antibody BAM705), mouse IL-13 (monoclonal capture antibody MAB413 and polyclonal detection antibody BAF413), human IL-5 (monoclonal capture antibody MAB405 and monoclonal detection antibody BAM6051), and human IL-13 (monoclonal capture antibody MAB213 and polyclonal detection antibody BAF213) were purchased from R&D Systems (Minneapolis, MN, USA).

For fluorescence flow cytometry of human ILC2s, FITC-conjugated human lineage antibody cocktails [CD3 (UCHT1), CD14 (HCD14), CD16 (3G8), CD19 (HIB19), CD20 (2H7), CD56 (HCD56)], PerCP/Cy5.5-conjugated anti-CD45 (HI30), PE-conjugated anti-CD127 (A019D5), and PE/Cy7-conjugated CD161 (HP-3G10) antibodies were obtained from BioLegend (San Diego, CA, USA), while the Alexa Fluor 647-conjugated CRTH2 (BM16) antibodies were obtained from BD Biosciences (Franklin Lakes, NJ, USA).

Lipidure BL802, a water-soluble polymer of 2-methacryloyloxy ethyl phosphorylcholine, was purchased from the NOF Corporation (Tokyo, Japan). Dimethyl pimelimidate-2HCl (21666) was purchased from Thermo Fisher Scientific (Rockford, IL, USA). CF660R streptavidin was purchased from Biotium (29040; Hayward, CA, USA). Cy3 streptavidin was purchased from Thermo Fisher Scientific (438315; Waltham, MA, USA).

The RPMI-1640 medium (R8758, Sigma-Aldrich, St. Louis, MO, USA) for mouse ILC2s and human peripheral ILC2s was supplemented with MEM nonessential amino acid solution (M7145, Sigma-Aldrich), 10 mM HEPES buffer (H3537, Sigma-Aldrich), 1 mM sodium pyruvate (11360-070, Gibco / Thermo Fisher Scientific), penicillin / streptomycin (15140-122, Gibco / Thermo Fisher Scientific), 55 µM 2-mercaptoethanol (21985-023, Gibco / Thermo Fisher Scientific), and 10 % FCS (S1820, Japan Bioserum, Tokyo, Japan). The medium was supplemented with 10 ng/ml recombinant mouse IL-2 (rmIL-2) (402-ML, R&D systems) for long-term culture of mouse ILC2s.

Stimuli for the mouse ILC2s were rmIL-2, rmIL-25, and rmIL-33 (402-ML, 1399-IL, and 3626-IL, R&D Systems). Stimuli for human ILC2s were recombinant human IL-2 (rhIL-2) (Imunace35, Shionogi, Osaka, Japan.), rhIL-25, rhIL-33, and rhTSLP (1258-IL/CF, 3625-IL/CF, and 1398-TS/CF, R&D Systems).

### Preparation of mouse ILC2

Mouse ILC2s were isolated from the mesenteric fat tissue of wild-type C57BL/6 female mice, as previously described^46^. Isolated mILC2s were cultured in conditioning medium with rmIL-2 (10 ng/ml) for several weeks and used for experiments.

### Isolation of ILC2s from human peripheral blood

Peripheral blood was obtained from healthy volunteers at Keio University School of Medicine as described previously^43^. This study was approved by the Institutional Review Board of the Keio University School of Medicine (approval number: 20090009). All participants provided written informed consent. Peripheral blood mononuclear cells (PBMCs) were separated using the Lymphoprep™ lymphocyte separation medium (Axis-Shield, Dundee, UK) according to the manufacturer’s protocols. PBMCs were stained with human lineage antibody cocktails (Lin) and anti-CD45, anti-CD127, anti-CRTH2, and anti-CD161 antibodies for 30 min at 2–8 °C. ILC2s were sorted as PI-Lin-CD45+CD127+CRTH2+CD161+ cells via flow cytometry using MoFlo™ XDP (Beckman Coulter, Brea, CA, USA).

### Optical arrangement

All measurements were performed with a completely automated inverted microscope (ECLIPSE Ti-E; Nikon, Tokyo, Japan) equipped with a high NA 60× objective lens (TIRF 60 × H; NA, 1.49; Nikon). The microscope was customised with the installation of white-light TIRF optics (high-performance Epi-fl illuminator module# TI-SFL) with LED light (X-Cite® XLED1, mounted with BGX:505-545 nm and RLX:615-655 nm; Excelitas Technologies Corp., Waltham, MA, USA). The laser light source used for comparison with the LED was purchased from Coherent (CUBE 640-40C; Santa Clara, CA, USA). Excitation filters (FF01-530/43 for Cy3 and FF01-635/18 for CF660R), emission filters (FF01-593/40 for Cy3 and FF01-692/40 for CF660R), and a dichroic mirror (FF560/659-Di01) were used. These optical filters were purchased from Semrock (Rochester, NY, USA). A semi-circular grinding acrylic lens block for measuring the TIR incident angle was purchased from KENIS Ltd. (H-100, refractive index 1.52; Osaka, Japan). Each image was projected onto a scientific CMOS camera (ORCA-Flash4.0 V2; Hamamatsu Photonics K.K., Shizuoka, Japan). A stage-top incubator (INUBG2TF-WSKM; Tokai Hit Co., Shizuoka, Japan) was used to control the temperature, humidity, and gas concentration. Control of the entire observation was performed using Nis-Elements AR 4.6 (Nikon).

### Designed for stable optical system

We focused on the light source for TIRFM to establish a long-term stable observation platform because the coherent laser light (commonly used for TIRFM) generated an optical interference pattern variation that was the main cause of measurement noise in the multi-point time-lapse observations, as the stage movement caused a slight shift in the optical path. One limitation of this method is that the directivity of the LED light is lower than that of the laser. To establish the total internal reflection preventing the leakage of propagating light to the sample area, the minimum angle of the incident LED light must be greater than the critical angle at the interface of the glass and materials of the well microstructure. For a microstructure on a glass surface fabricated with PDMS, which is one of the most used materials for microfabricated chips employed in biological applications, the critical angle at the interface between the PDMS and glass is 67.1° as calculated by sin^-1^(n_PDMS_/n_Glass_). Thus, the light incident at the critical angle on the interface between the aqueous solution and glass led to stray light generation in PDMS and high levels of background light (Fig. 1c, d); therefore, the excitation light should be incident at a larger angle. In this case, the light from the LED with low directivity was blocked by the aperture limit of the objective lens (sin^-1^(N_Aobj._/n_Glass_) = 78.6°), resulting in insufficient excitation intensity (Fig. 1d). To overcome this issue using innovations in microfabrication structures, we developed a TIRF-dish in which a refractive index adjustment layer of an amorphous fluoropolymer (AF layer; nAF = 1.34) was inserted at the interface between the PDMS and glass. In the TIRF-dish, the critical angle for the total reflection was found to be sufficiently small (sin^-1^(n_AF_/n_Glass_) = 61.8°), thereby greatly improving the signal intensity and the signal-to-background ratio (Fig. 1c-f).

### TIRF-dish fabrication

The TIRF-dish was composed of PDMS (Sylgard184; Dow Corning Toray, Tokyo, Japan), microscopic-grade coverslips (25 × 25 mm No. 1 cat# C025251, Matsunami Glass Ind., Osaka, Japan), and an amorphous fluoropolymer (CYTOP; AGC Chemicals Company, Tokyo, Japan), which served as the reflection index matching layer with the aqueous solution. The TIRF-dish was prepared as follows: a mould pattern, comprising 996 arrays that were 80 μm in diameter or had square through-holes with 115 μm centre-to-centre spacing and 80 μm thickness, were fabricated on a Si wafer with SU-8 3050 (NIPPON KAYAKU, Tokyo, Japan) according to the manufacturer’s instructions. PDMS (base: curing agent = 10:1) was poured onto the SU-8 mould and covered with a surface-inactivated glass slide. Then, the mould-PDMS-glass slide complex was pressed and cured at 80 °C for 1 h to form a through-hole array. The chamber block of the TIRF-dish was made with PDMS using a 3D fabricated mould (PEEK450G, Yasojima Proceed Co., Ltd., Osaka, Japan). The PDMS sheets and chamber blocks were immersed in n-hexane to remove any inhibitors of cellular activity. The cleaned PDMS sheet and chamber block were dried and permanently bonded using air plasma (SEDE-PFA; Meiwafosis Co., Ltd., Tokyo, Japan). The coverslip was cleaned by sonication in 5 N KOH for 1 h. The surface of the coverslip was aminated with VECTABOND reagent (SP-1800; Vector Laboratories, Burlingame, CA, USA), spin-coated with CYTOP 809A at 1,500 rpm for 20 s and cured at 80 °C for 1 h. Then the PDMS sheet with the chamber block was permanently bound to the surface-modified CYTOP-coated coverslip and baked at 105 °C for 3 h. The CYTOP coating exposed at the bottom of each nanolitre well was removed by wet etching using the CYTOP solvent (CT-solv100E and CT-solv180; AGC Chemicals Company) and cleaned by air plasma to expose the glass surface. Then, the bottom glass surface was aminated using the VECTABOND reagent. A mixture of capture antibodies (final concentration of 100 μg/ml) and dimethyl pimelimidate-2HCl (final concentration of 7 mg/ml) was loaded into each chamber to fix the capture antibodies on the bottom surface of each nanolitre well. The remaining reaction groups were blocked with monoethanolamine (0.1 M, pH 8.2). The TIRF-dish with immobilised antibodies was stored in phosphate-buffered saline supplemented with Lipidure BL802 reagent (0.2 % w/v) at 4 °C until further use.

### Preparation of the detection medium

The biotin-labelled detection antibody was coupled with either CF660R-labelled streptavidin or Cy3-labelled streptavidin at 1:10 molar ratios in the dark for 3 h. Unoccupied sites on streptavidin were blocked with excess Dpeg4-biotin acid (10199; Quanta BioDesign, Ltd., Powell, OH, USA). Unconjugated streptavidin and Dpeg4-biotin were removed via ultrafiltration (Amicon Ultra-0.5, 100 kDa; Merck Millipore, Billerica, MA, USA). The detection media contained the prepared CF660R and Cy3-labelled detection antibodies for each antigen with final concentrations as follows: anti-mIL-5 (90 nM for Fig. 1 and Supplementary Video 1, 30 nM for others), anti-mIL-13 (30 nM), anti-hIL-5 (22.5 nM), anti-hIL-13 (90 nM), BSA (1 % w/v), and the indicated combinations of the following additives (at final concentrations): rhIL-2 (20 U/ml), rhIL-25 (50 ng/ml), rhIL-33 (50 ng/ml), rhTSLP (50 ng/ml), rmIL-2 (10 ng/ml), rmIL-25 (10 ng/ml), and rmIL-33 (10 ng/ml).

### Observation of type 2 response from individual ILC2s

An aliquot of 500 cultured mouse ILC2s or 100 to 300 of freshly isolated human ILC2s was introduced into a chamber on a TIRF-dish and deposited in nanolitre wells via centrifugation (200 × *g* for 30 s). Accordingly, each nanolitre well contained zero to several mILC2s (bright field in Figure 3b), and approximately 200–300 nanolitre wells containing single mILC2s were used for statistical analysis. In the hILC2 experiment, which was designed to detect rare cellular activity, all wells containing hILC2s were tracked during the 3-day monitoring period, and statistical analysis was performed using the total number of hILC2s at the initial time frame. The intensity values of the empty wells were used as references to compensate for the spatiotemporal changes in the excitation light intensity and to calculate the dynamic threshold. The culture supernatant was replaced with freshly prepared detection medium immediately before observation. Mineral oil (M8410; Sigma-Aldrich) was layered on top of the detection medium to prevent evaporation. All wells in the TIRF-dish (996 wells × 4 chambers) were scanned using approximately 1-h cycles for 1 d or more. During the long-term tracking of hILC2s, data within 72 h were used for statistical analysis for the following two reasons. First, there was a concern that the overproduction of cytokines following repeated division would be underestimated by the quantification due to the limited capacity of the capture antibody. Second, hILC2s that showed repeated divisions were observed to overflow from a well and migrate to neighbouring wells.

### Image processing of the DSS for qLCI-S with high spatiotemporal resolution

The signal of the time-resolved fluorescence immunoassay in the LCI-S measurement environment can theoretically be regarded as a convolution of the temporal change in the concentration of the captured antigen and the association-dissociation kinetics of the fluorescent antibody (Supplementary Fig. 4). Therefore, we were able to computationally deconvolute the secretion activities from the detected fluorescence signal by referring to the staining kinetics obtained in the calibration measurements (Supplementary Fig. 5)^20^. First, a sequence of images of the same well was aligned temporally with sub-pixel accuracy. Next, a moving median filter for every five frames was applied to all pixels, followed by binning of the 5 × 5 pixels. Then, temporal deconvolution was performed on each pixel with reference to a function fitted to the antibody staining kinetics. Note that if a value fell below zero during the deconvolution process, it was considered noise, and the value was rounded up to zero at that time. This process suppressed the divergence of the accumulated computational noise.

### Image analysis for the statistical comparison of high-throughput qLCI-S measurement

The secretion activity of the cells in each well was analysed using an in-house VBA programme in Microsoft Excel using the mean intensities and cell numbers of each well at each time frame. The mean intensity of each well was measured from IL-5 and IL-13 secretion signal images using commercially available image analysis software (NIS-elements AR 4.6, Nikon). The number of cells in each well was counted manually or estimated by an in-house artificial intelligence algorithm^47^. First, we corrected the intensity deviation between the wells for the field-of-view configuration and the time period. The mean fluorescence intensities of empty wells were summarised and averaged for each configuration (upper left, upper right, lower left, and lower right in the field of view), each chamber, and each time frame. The quotients of these values to the total average were calculated as coefficients, and the intensities of the individual wells were corrected using these coefficients. Next, we subtracted the initial intensity from the intensity at each time frame for each well. For wells with an initial intensity outlier larger than 1.5 × (interquartile range), the average intensity of empty wells was subtracted. This calculation eliminates the contribution of autofluorescence from infrequent contaminants and well walls in each well. Third, the influence of antibody staining kinetics and cross-staining of IL-5 and IL-13 was removed by temporal deconvolution. The calculated values were filtered by temporal integration and moving the median to reduce the processing noise, which were then used for statistical evaluation of the secretion activity of individual cells.

The detection of secretion positivity for the intensity of each well was performed using a detection limit [3 × standard deviation (SD)] for the intensity of the empty wells. This SD, however, was estimated using the median absolute deviation of the empty wells multiplied by 1.4826. Additionally, taking advantage of the continuity of the accumulating signal in the temporal direction, we judged a secretion signal trajectory to be positive if it was above the threshold of ≥ 2 time frames, while allowing for a transient fall below the threshold if it was within 3 time frames. The determination of hyperactive hILC2 was performed in conjunction with the above secretion positivity using a threshold of 50× SD corresponding to the results of the k-means cluster analysis (Fig. 4 g).

### Statistical analysis

The secretion signals were statistically analysed using the Mann–Whitney test or log-rank test. The false discovery rate of each test was controlled using the Benjamini–Hochberg procedure. Statistical significance was set at P < 0.05. These graphs were drawn, and the tests were performed using the statistical software OriginPro (Origin Lab, Northampton, MA, USA).

### Functional RNA-Seq on LCI-S

ILC2s were retrieved from the TIRF-dish using a glass capillary (L-Tip 15 mm 60° 15 mm, Yodaka Co., Ltd., Kanagawa, Japan) controlled by a pneumatic microinjector (IM-11-2, Narishige, Tokyo, Japan) and a micromanipulator (Transferman NK2, Eppendorf, Hamburg, Germany) and transferred into 4 ml RNase-free water (06442-95, Nacalai Tesque, Kyoto, Japan) in 0.2-Ml polymerase chain reaction tubes. Isolated cells were spun down, flash frozen in liquid nitrogen, and stored at −80 °C until further use. We obtained cDNA from poly(A) RNA of each sample using a SMART-Seq v4 Ultra Low Input RNA Kit for Sequencing (Takara Bio Inc., Shiga, Japan), following the manufacturer’s instructions. The pre-amplified cDNA was obtained after reverse transcription (90 min at 42 °C, 10 min at 70 °C) and pre-amplification (24 cycles (N1–N4) or 18 cycles (P1–P3): 1 min at 95 °C, 10 s at 98 °C, 30 s at 65 °C, and 3 min at 68 °C). After the purification of the amplified cDNA using Agencourt AMPure XP beads (A63881, Beckman Coulter), cDNA sequences were analysed using Illumina HiSeq with an entrusted analysis service (50 bp single-read, Medical & Biological Laboratories Co., Ltd. Japan). The obtained sequence data were mapped to the reference genome (GRCh38.87) using Bowtie2 (version 2.3.1), and the number of transcripts was counted using the RSEM software package (1.3.0). Subsequent Reproducibility-Optimised Test Statistic (ROTS) analysis of the expressed genes with TPM ≥ 1 from one or more samples detected 1,869 differentially expressed genes (DEGs). The distribution of the TPMs calculated from the count data for each cell showed a continuous distribution in hyperactive ILC2, whereas the distribution obtained from silent ILC2 showed a sharp decrease in the number of genes with TPMs below 50 (Supplementary Fig. 6). This phenomenon is seen when cDNA amplification is performed using a low absolute amount of transcripts and caused by limits in the efficiency of reverse transcription and amplification. We found no significant influence of the cells recovered from the qLCI-S platform, allowing us to obtain data comparable to normal single-cell RNA-Seq. The detection threshold was set at 44.7 (Supplementary Fig. 6), and significant difference analysis was performed for genes with TPM values above the threshold in the hyperactive ILC2s.

## Acknowledgements

The authors would like to thank Mr. Kazuki Yoda for his scientific discussion. This manuscript was edited by Editage.

## Funding

This study was supported by JST PRESTO (grant no. JP17940748 to Y.S.), AMED (grant no. JP18ek0410031, JP20hm0102082), the ImPACT Program of the Council for Science, Technology, and Innovation (Cabinet Office, Government of Japan), the Japan Society for the Promotion of Science (grant no. 20H04512, Core-to-Core Program), and White Rock Foundation.

## Author Contributions

M.Y. and Y.S. conceived the original idea and designed the experiments. M.Y., K.M., Y.T., N.S., and Y.S. established the measurement equipment and environment and the analysis environment. M.Y., K.M., T.K., and Y.S. performed and analysed the experiments. Y.M., T.K. S.K., and K.M. provided the murine sample. T.K., H.K., R.B., M.M., and K.F. prepared the human sample. M.Y., K.M., T.K., H.K., R.B., M.M., Y.M., T.K., S.K., K.F., K.M., S.U., and Y.S. interpreted the biological significance of the results. M.Y., K.M., T.K., T.F., and Y.S. examined the statistical adequacy of the analysis. M.Y., K.M., K.M., S.U., and Y.S. interpreted the data and wrote the paper. K.G., O.O., and T.F. helped supervise the project. K.F., K.M., S.U., and Y.S. supervised the project.

## Competing Interests

The University of Tokyo has filed a patent application covering the methodology of the cell recovery system described in this paper. M.Y. has a financial interest in Live Cell Diagnosis, Ltd., which is a company that was launched for the development of LCI-S. N.Y. is currently employed by Live Cell Diagnosis. K.G. is a shareholder of CYBO and Cupido.

## Data availability

Datasets generated and/or analysed during the current study are available from the corresponding authors on reasonable request.

## Extended data

**Supplementary Figure 1.**
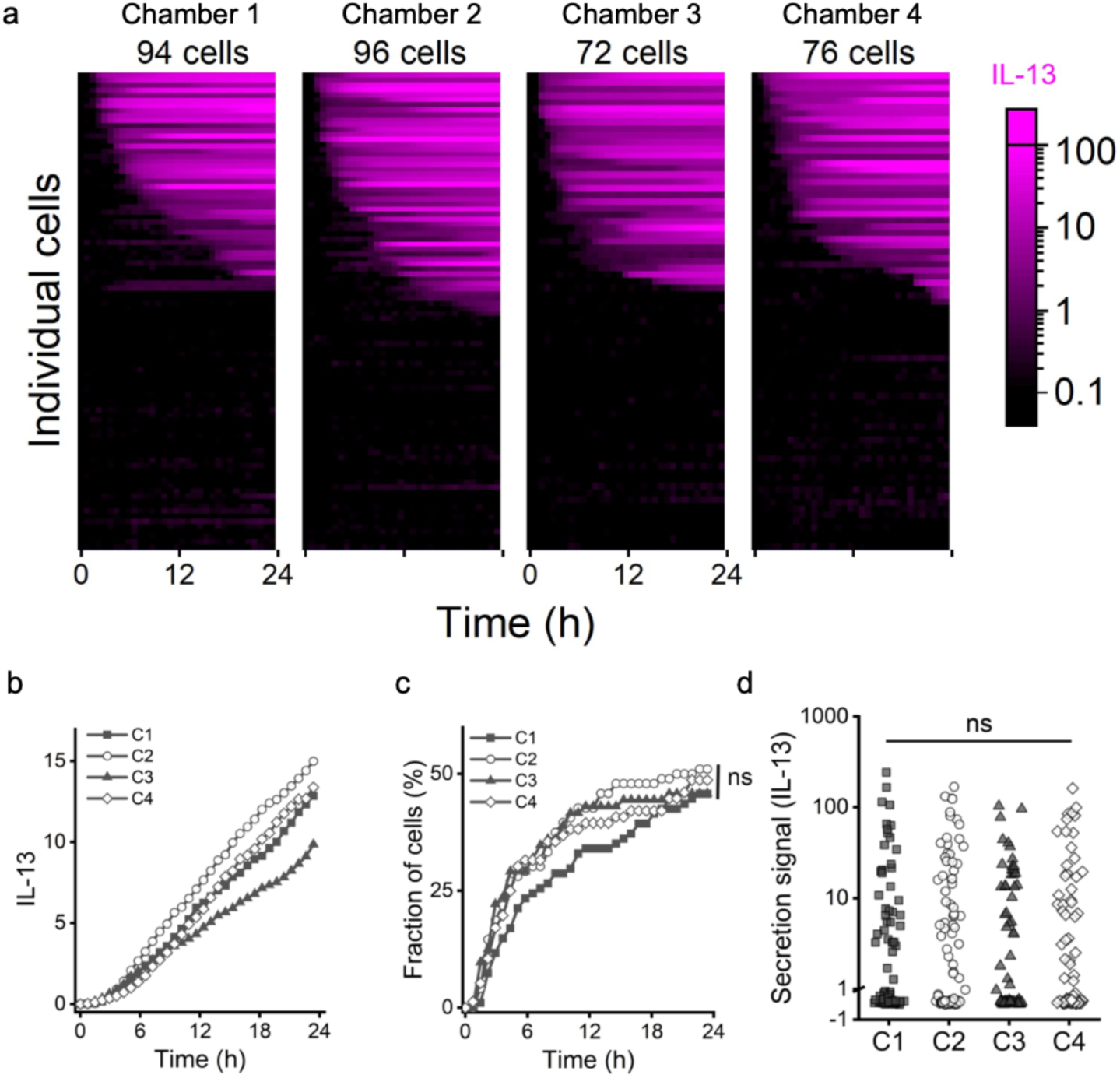
Evaluation of the reproducibility and independence of the four chambers of the TIRF-dish. **(a)** Chronological table of the cumulative secretion signals of every single cell arranged for each sample chamber (C1–C4) with the same culture environment. Each row reflects the dynamic activity of an individual cell. Colour scale bars show the intensity of IL-13 secretion signals. **(b)** Averaged cumulative secretion signal of IL-13 in each sample chamber. **(c)** Proportion of secretion-positive cells in each sample chamber. Differences were tested using the log-rank test. **(d)** Maximum cumulative secretion signal in each sample chamber. Differences were assessed using the Mann-Whitney *U* test.

**Supplementary Figure 2.**
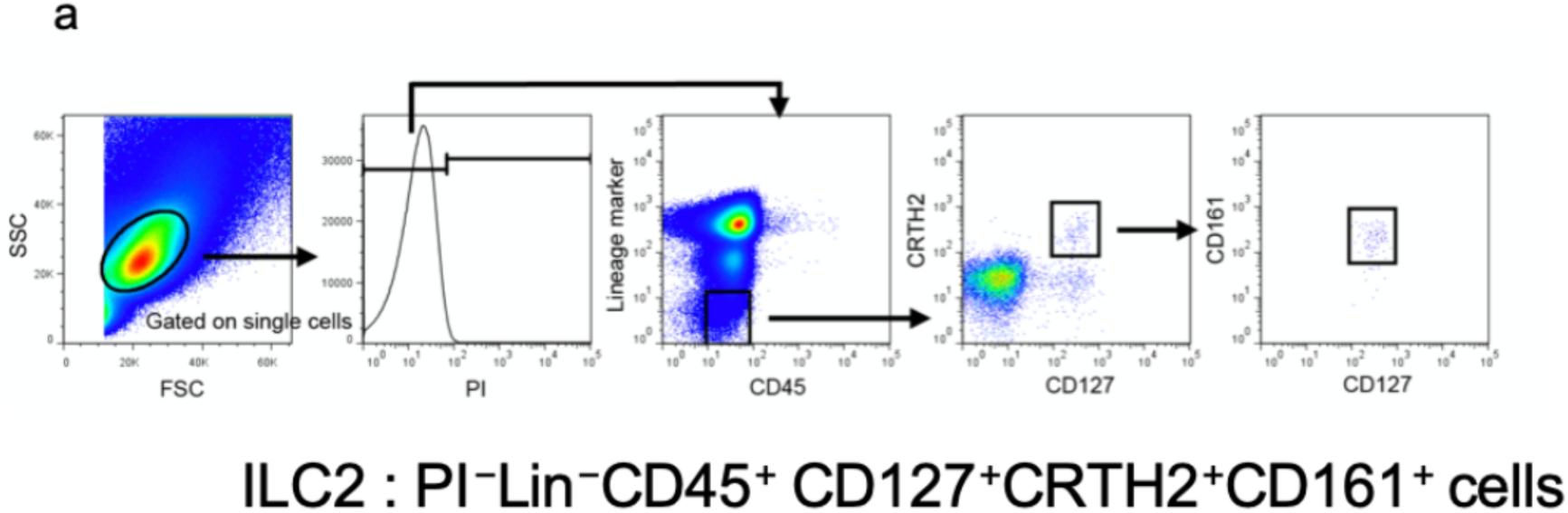
Purification of human peripheral ILC2s. Fluorescence flow cytometry plots showing gating strategy for sorting human ILC2s. Peripheral blood mononuclear cells were isolated using Lymphoprep and labelled with several surface differentiation markers, as described in the Materials and Methods. The labelled cells were sorted via fluorescence-assisted cell sorting (FACS) and identified as ILC2s.

**Supplementary Figure 3.**
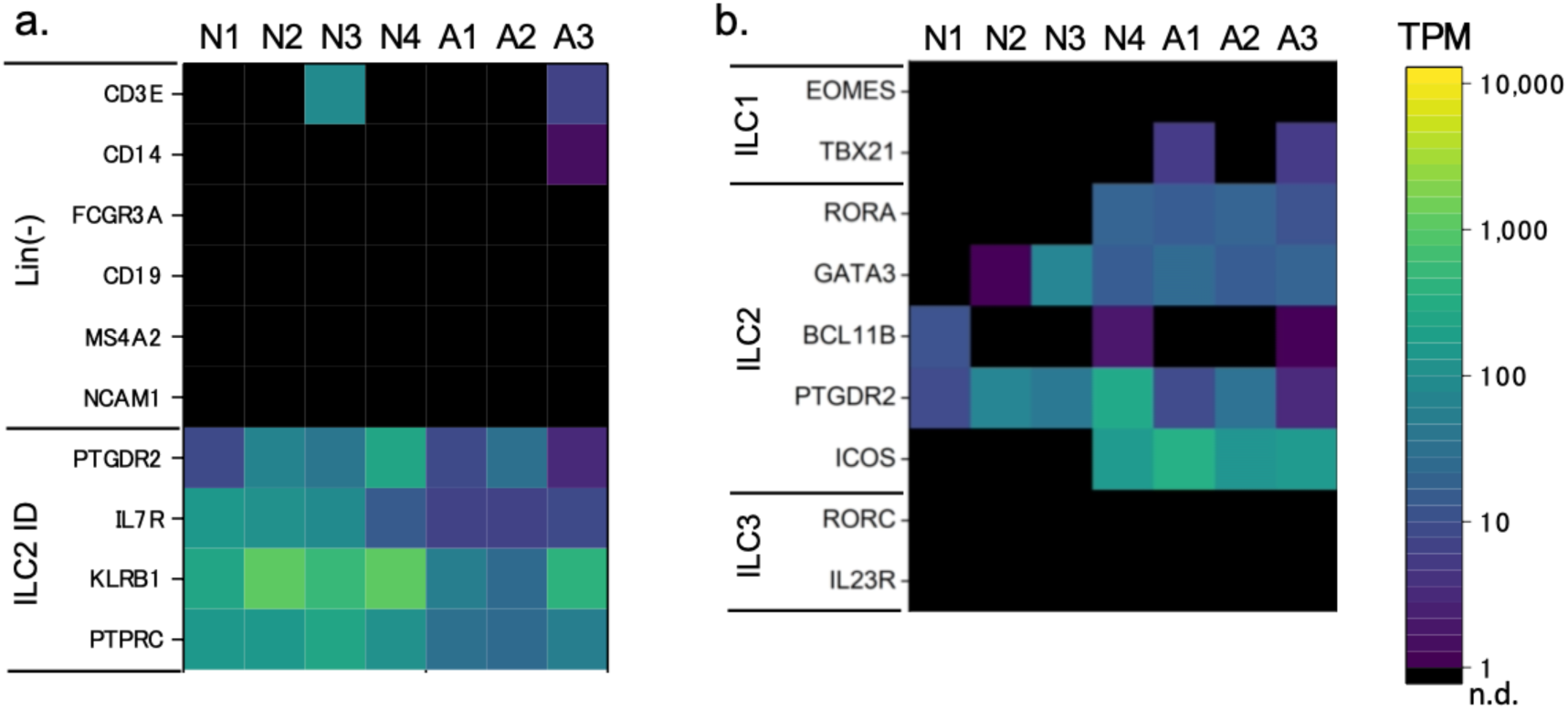
Expression differences of active and silent ILC2s. **(a)** Transcript expression of genes used for FACS purification showed in parallel to proteins. Lineage markers were rarely detected from both active and silent populations. In contrast, identity markers for ILC2 were not only detected from active populations but also specifically detected from silent populations exceeding the confidence level. **(b)** Comparisons of the expression of specific genes of each ILC type. ILC2-specific genes were detected in both silent and active populations, except for BCL11B. Genes specific for ILC1 and ILC3 were rarely detected.

**Supplementary Figure 4.**
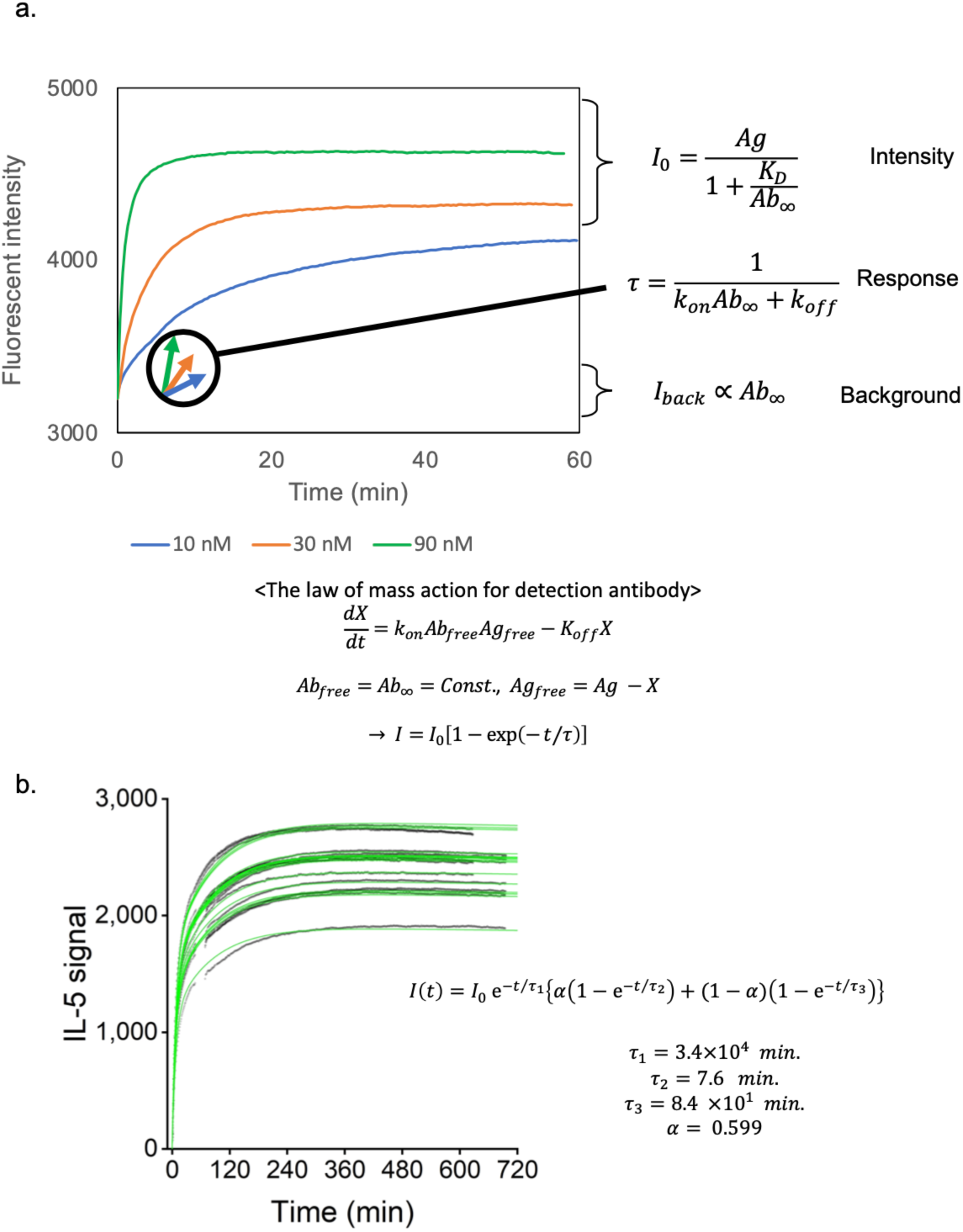
Binding dynamics of detection antibody. The typical binding dynamics of the fluorescent sandwich immunocomplex are shown. In this assay, transient secretion activity was mimicked by the instantaneous release of a recombinant protein by microinjection. A high local concentration of the capture antibody enables the system to immediately immobilise the secreted antigen compared to that in the measurement interval of the time-lapse. Immediate immobilisation is also reflected in the localised pattern of the secretion signal. Compared with the fast dynamics of capturing, the staining dynamics using the detection antibody require time to reach equilibrium. The law of mass action for single-step reactions explains the dynamics of the time-resolved fluorescence immunoassay used in LCI-S. **(a)** Demonstration of a time-resolved fluorescence immunoassay for human IL-5. IL-5 (100 ng/mL) was captured on an anti-human IL-5 antibody-coated glass bottom. After washout of excess unbound IL-5, TIRF measurements were performed. The intensities increased immediately after the addition of detection antibodies at a final concentration of 10, 30, or 90 nM. The maximal intensity was controlled by the efficiency of forming an immunocomplex at equilibrium, depending on the amount of antigen and the affinity and concentration of the detection antibody. The response speed was controlled by the affinity and concentration of the detection antibodies. The background intensity is proportional to the concentration of the detection antibody, and the detection sensitivity generally decreases in proportion to the square root of the background light intensity. The cost of measurement increased in proportion to the concentration of the detection antibody. **(b)** Time-resolved fluorescence immunoassay every 1 min for recombinant mouse IL-5 in different amounts microinjected into a nanolitre well in a TIRF-dish with 90 nM of anti-mouse IL-5 detection antibody. The curves of the formula indicated on the graph fit well with the data. The first term with τ_1_ indicates the exponential decay of the secretion signal; meanwhile, the second term with τ_2_ and the third term with τ_3_ indicates the fast and slow exponential growth of the secretion signal with ratio α, respectively. The fitted parameter was used to calculate the secretion activity (Fig. 1g-i).

**Supplementary Figure 5.**
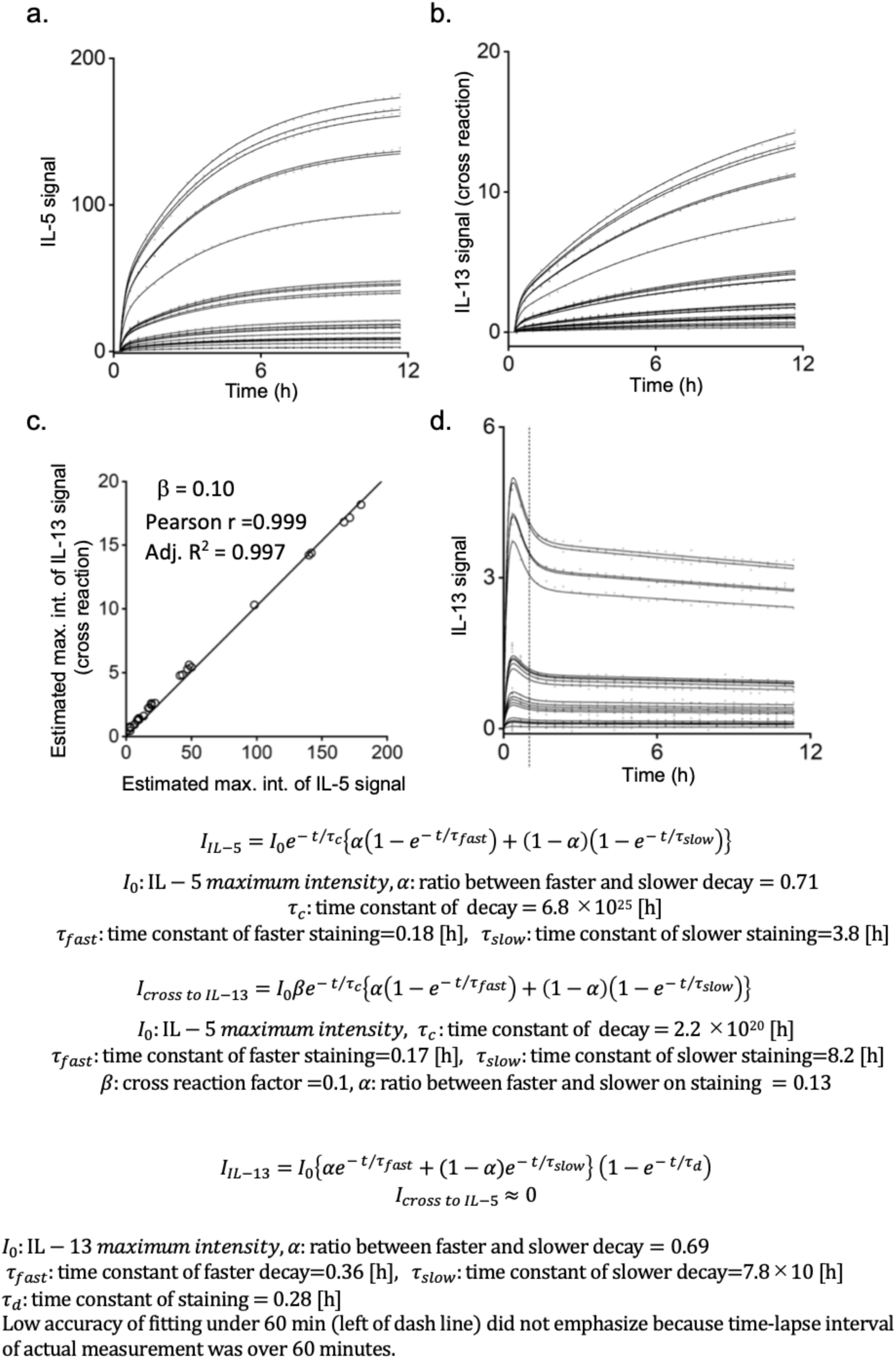
Standard curve of fluorescent immunoassay signals of recombinant human IL-5 or IL-13 sampled by micro-injecting proteins. **(a)** Time-resolved fluorescence immunoassay for recombinant human IL-5 microinjected into nanolitre wells in a TIRF-dish at different amounts performed with 22.5 nM of anti-mouse IL-5 detection antibody every 20 min. **(b)** Time-resolved fluorescence immunoassay for recombinant human IL-5 microinjected into nanolitre wells in a TIRF-dish at different amounts performed with 90 nM of anti-human IL-13 detection antibody every 20 min. The curves of the formula for *IIL-5* and *Icross to IL-13* fitted well with the data shown in panel a and b, respectively. **(c)** Kinetics of cross-reactivity of human IL-13 protein to human IL-5 fluorescence immunoassay was measured by depositing trace amounts of recombinant human IL-13 into a TIRF-dish carrying an immunoassay constructed with the human IL-5 antibody set. Each dot indicates the maximum intensity estimated by the formulas for *I_IL-5_* and *I_cross to IL-13_*. The human anti-IL-13 antibody had cross-reactivity with the IL-5 sandwich immunocomplex, whereas the anti-IL-5 antibody had no cross-reactivity with the IL-13 sandwich immunocomplex. **(d)** Time-resolved fluoroimmunoassay for recombinant human IL-13 microinjected into nanolitre wells in a TIRF-dish at different amounts was observed with 90 nM of anti-human IL-13 detection antibody every 20 min. The fitted parameters were used in Figures 4a and 4b.

**Supplementary Figure 6.**
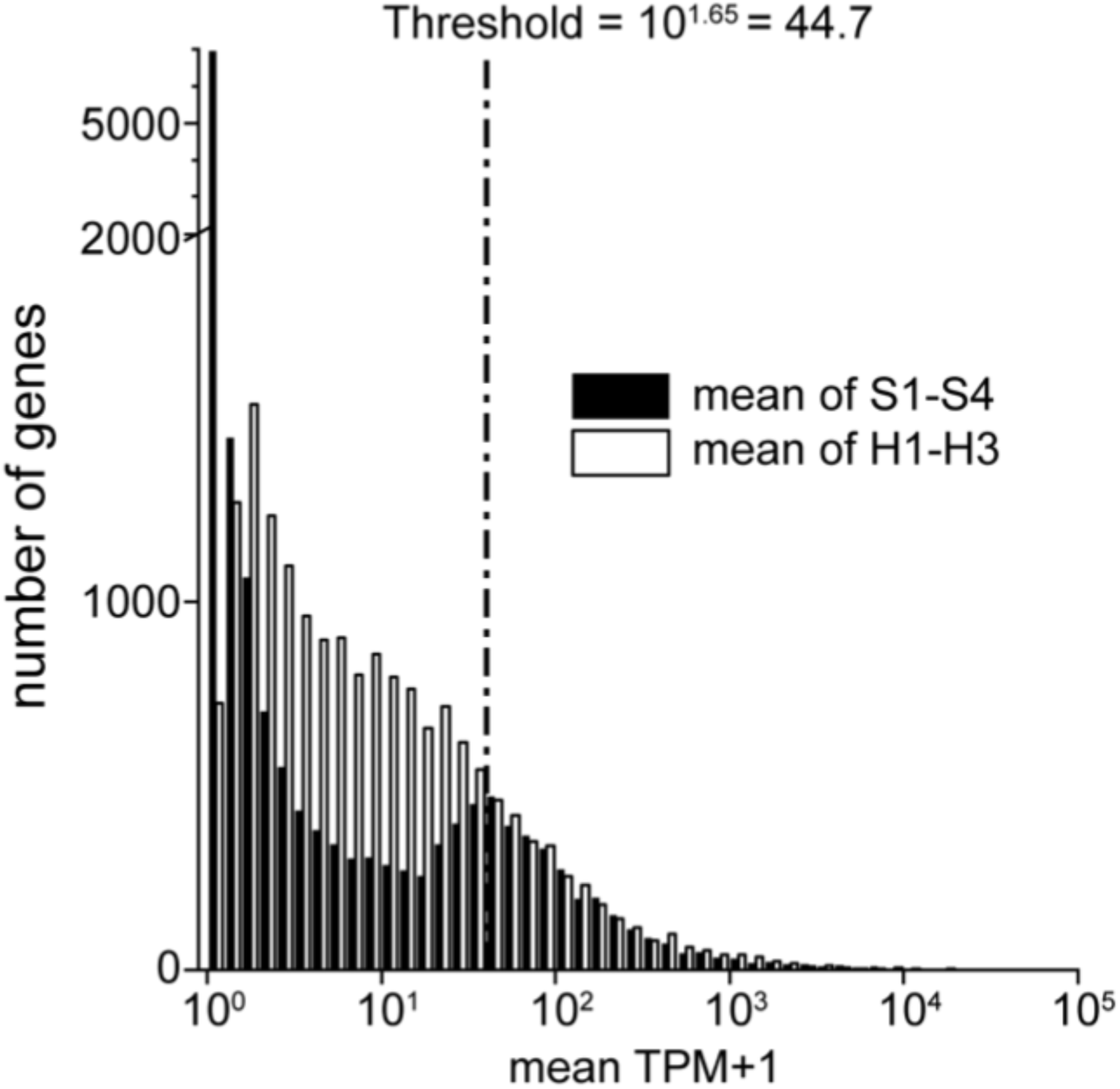
Distribution of transcripts per kilobase million (TPM) for retrieved hILC2s. Mean TPM of hyperactive or silent hILC2s. The values were increased by 1 to show all genes in a logarithmic manner. Silent hILC2s (S1 to S4), whose cDNA library was derived from a single cell, showed a cut-off pattern in comparison with multi proliferated hyperactive hILC2s. We defined the confidence level associated with stochastically missing gene expression as TPM = 43.7. We assumed that the difference in gene expression was more reliable if the expression level of the gene exceeds this threshold, especially in silent hILC2s.

**Supplementary video 1|** Visualisation of dynamic IL-5 secretion activity of mILC2. Bright-field images of a mouse innate lymphoid cell (mILC2; upper left), cumulative secretion signal (CSS) images of IL-5 (lower left), and temporally deconvoluted secretion signal (DSS) images (lower right). The intensities of CSS and DSS are in pseudo-colour, as shown on the colour bar of each panel. The white outline in each panel indicates the shape of the cell in the bright-field image. The interval of each image is 1 min.

**Supplementary video 2|** Visualisation of dynamic IL-5/IL-13 secretion activity of hILC2. Bright-field images of innate lymphoid cells from human peripheral blood (hILC2s; upper), cumulative secretion signal (CSS) images of IL-5 and IL-13 (bottom), and temporally deconvoluted secretion signal (DSS) images (middle). The intensities of the secretion signals of IL-5 and IL-13 are shown as green and magenta pseudo-colours, respectively.The interval of each image is 1 h.

